# Temporal patterns in the evolutionary genetic distance of SARS-CoV-2 during the COVID-19 pandemic

**DOI:** 10.1101/2020.11.01.363739

**Authors:** Jingzhi Lou, Shi Zhao, Lirong Cao, Zigui Chen, Renee WY Chan, Marc KC Chong, Benny CY Zee, Paul KS Chan, Maggie H Wang

## Abstract

**Background:** During the pandemic of coronavirus disease 2019 (COVID-19), the genetic mutations occurred in severe acute respiratory syndrome coronavirus 2 (SARS-CoV-2) cumulatively or sporadically. In this study, we employed a computational approach to identify and trace the emerging patterns of the SARS-CoV-2 mutations, and quantify accumulative genetic distance across different periods and proteins.

**Methods:** Full-length human SARS-CoV-2 strains in United Kingdom were collected. We investigated the temporal variation in the evolutionary genetic distance defined by the Hamming distance since the start of COVID-19 pandemic.

**Findings:** Our results showed that the SARS-CoV-2 was in the process of continuous evolution, mainly involved in spike protein (S protein), the RNA-dependent RNA polymerase (RdRp) region of open reading frame 1 (ORF1) and nucleocapsid protein (N protein). By contrast, mutations in other proteins were sporadic and genetic distance to the initial sequenced strain did not show an increasing trend.

## Introduction

The pandemic of coronavirus disease 2019 (COVID-19) caused by the severe acute respiratory syndrome coronavirus 2 (SARS-CoV-2) poses severe threat to public health globally. The genetic mutations in SARS-CoV-2 have been detected frequently. Although most mutations occur sporadically and are purged shortly, it appears that some mutations gradually reach fixation [1]. Given increasing numbers of fixed mutations, the circulating SARS-CoV-2 strains may diverge from the original strain in terms of an increasing accumulative genetic distance. The accumulation of genetic distance reflects a steady viral evolutionary process that may affect the characteristics of SARS-CoV-2, immune recognition, and effectiveness of antivirals targeting the pathogen. In this study, we employed a computational approach to identify and trace the emerging patterns of the SARS-CoV-2 mutations, and quantify accumulative genetic distance across different periods and proteins.

## Data and methods

The full-length human SARS-CoV-2 strains in United Kingdom were obtained from Global Initiative on Sharing all Influenza Data (GISAID) [2] on September 23, 2020. A total of 40,527 strains were collected with the collection date ranging from January 27 to September 14, 2020. We used a stratified sampling scheme to randomly selected sequences in biweekly time interval. See Supplementary Materials S1 for the details of the sampling scheme and summary. As one of the first reported sequences, ‘China/Wuhan-Hu-1/2019’, which was collected in December 31, 2019, was considered as the reference strain for sequence alignment, and as the initial strain for genetic distance calculation against other strains. The genetic distance to the initial strain was defined by the Hamming distance. We investigated the temporal variation in the accumulative genetic distance in the cross-sectional series. To better observe the dynamic trend of genetic distance, a sliding window was applied (Supplementary Materials S2). Multiple sequences alignment was performed using MEGA-X (version 10.1.8).

## Results and discussions

We observed that the genetic distance to the initial strain steadily increased with time since February 2020 (Figure 1). The overall distance of all proteins rose from 0 to a peaking level at 9.27 codons on August 21, 2020, which implied mutations with evolutionary advantages occurred and gradually accumulated. Moreover, deleterious mutants can only preserve for a short period in virus population and disappeared due to functional issues or less adaption to environment, which explains the fluctuations in the genetic distance curve, e.g. open reading frame 8 (ORF8) in Figure 1. By further observing the distance in each protein, we found continuous increasing trends in the spike protein (S protein), the ORF1 (especially in the RNA-dependent RNA polymerase, Supplementary Materials S3) and the nucleocapsid protein (N protein), and no obvious trends in the membrane protein (M protein), the envelope protein (E protein), the ORF3, ORF6-8 and ORF10, see Figure 1. It suggested a stronger natural selection pressure on the S protein, the ORF1 and the N protein. The S protein is responsible for receptor recognition and membrane fusion, and contains a receptor binding domain (RBD), which is considered as the target for the SARS-CoV-2 vaccines under development[3–5]. The RdRp in ORF1 plays a central role in the replication and transcription cycle of SARS-CoV-2 and thus is considered as a target for nucleotide analog antiviral inhibitors [6]. N protein is a multifunctional and highly immunogenic determinant, whose function is mediating the packaging of the viral RNA genome into the nascent virion[7]. These three proteins were critical in determining the course of transmission, infection and reproduction of the virus, and thus accumulated mutations in these proteins might be related to the viral adaptation to the environment both in vivo and in vitro. By contrast, genetic distances of the proteins with sporadic mutation were usually below 0.5 codon without demonstration of an increasing trend. To date, the mutations in these proteins have not maintained, which might be due to their less competitive strength in facilitating the viral adaptation to the environment.

**Figure 1.**
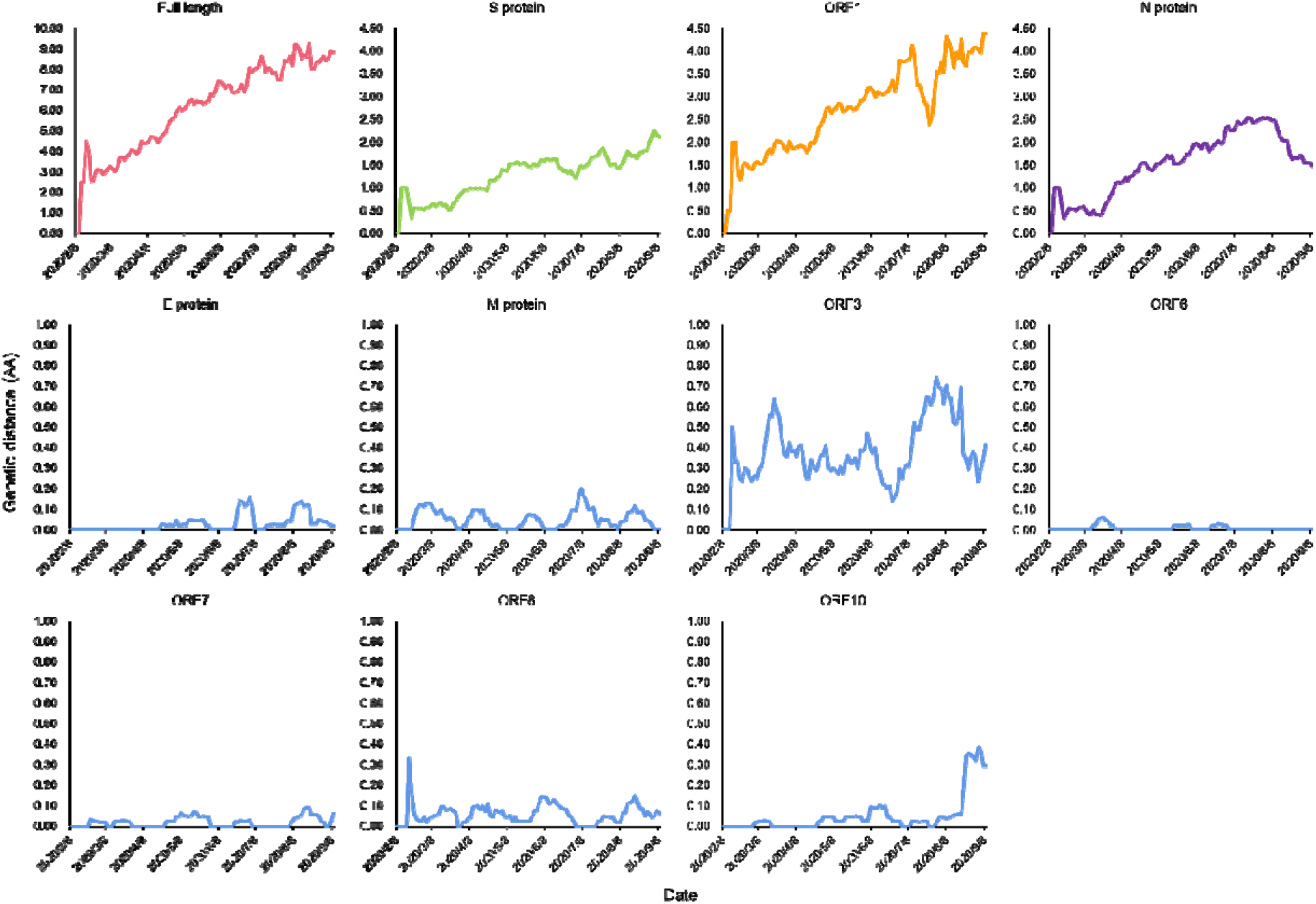
The time-varying genetic distance from initial strain in different proteins. The green, orange and purple lines represent genetic distance in S protein, ORF1 and N protein with increasing trends, respectively. The blue lines represent genetic distance in M protein, E protein, ORF3, ORF6, ORF7, ORF8 and ORF10 without notable trends.

Our results showed that the SARS-CoV-2 was in the process of continuous evolution, mainly involved in the S protein, the RdRp region of ORF1 and the N protein. In August 24, 2020, the first cases of COVID-19 re-infection was officially identified and reported that the viruses corresponding to the first and second infections carried 13 amino acid differences [8]. Another study showed that functional S protein and N-terminal domain of SARS-CoV-2 with mutations conferred resistance to monoclonal antibodies [9]. Continuous evolution of the virus might bring considerable challenge to the development of antiviral drugs and vaccines.

## Conclusion

This study presents the evolutionary process of SARS-CoV-2 virus from the aspect of genetic distance. The continuous mutation accumulation was observed in genes encoding the S protein, the RdRp region and the N protein, but not be observed in genes encoding other proteins to date. Therefore, future investigation is warranted to study the characteristics and the effects associated with the accumulative mutations in SARS-CoV-2, as well as the treatment or control strategies.

## Declarations

## Authors’ contributions

MHW conceived the study. JL collected the data and carried out the analysis. JL, SZ, LC and MHW discussed the results. JL drafted the first manuscript. JL, SZ, LC and MHW reviewed and edited the manuscript. All authors critically read and revised the manuscript and gave final approval for publication.

## Disclaimer

The funding agencies had no role in the design and conduct of the study; collection, management, analysis, and interpretation of the data; preparation, review, or approval of the manuscript; or decision to submit the manuscript for publication.

## Acknowledgements

The SARS-CoV-2 sequences were collected from Global Initiative on Sharing all Influenza Data (GISAID) accessible via https://www.gisaid.org/. We thank the contribution of the submitting and the originating laboratories. This study was conducted using the resources of Alibaba Cloud Intelligence High Performance Cluster computing facilities, which is made free for COVID-19 research.

## Ethics approval and consent to participate

The human SARS-CoV-2 strains were collected via public domains, and thus neither ethical approval nor individual consent was not applicable.

## Availability of materials

All data used in this work were publicly available.

