## Supplementary materials for "Temporal patterns in the evolutionary genetic distance of SARS-CoV-2 during the COVID-19 pandemic"

**Supplementary Information**

**Table of content**

### **S1. SARS-CoV-2 sequencing data and COVID-19 number of cases surveillance data**

The full-length human SARS-CoV-2 strains in United Kingdom were collected from Global Initiative on Sharing all Influenza Data (GISAID) [1] on September 23, 2020. There were 40,527 strains with the collection date ranging from January 27 to September 14, 2020. Table S1 summarized the number of human SARS-CoV-2 strains in GISAID and sample size used in this study. We set 22 successive periods with the same time interval. Especially, there was no further division in January and February, because the number of strains were limited and the disease was just beginning to spread at this stage in United Kingdom. Since the number of sample strains varied in different periods, we randomly sampled 30 strains in the period when there were more than 30 strains. This sampling scheme aimed to balance the weights due to different sample sizes that may affect the sliding window framework applied in calculating genetic distance, see section S2. A total of 618 complete strains were finally analyzed in this study.

The multiple sequence alignment was performed by MEGA-X (version 10.1.8) via Alibaba Cloud platform, and the strain China/Wuhan-Hu-1/2019|EPI_ISL_402125 was used as a reference strain. The protein translation was conducted by ***R*** (version 3.6.3).

Table S1. Number of human SARS-CoV-2 strains in GISAID and sample size included in this study for different periods.

| Period | Number of strains | |
| --- | --- | --- |
|  | In GISAID | In this study |
| Jan 1^st^ – 31^st^ | 3 | - |
| Feb 1^st^ – 29^th^ | 119 | 30 |
| Mar 1^st^ - 10^th^ | 972 | 30 |
| Mar 11^th^ - 20^th^ | 2,580 | 30 |
| Mar 21^st^ - 31^st^ | 7,471 | 30 |
| Apr 1^st^ - 10^th^ | 7,313 | 30 |
| Apr 11^th^ - 20^th^ | 5,727 | 30 |
| Apr 21^st^ - 30^th^ | 4,156 | 30 |
| May 1^st^ – 10^th^ | 2,524 | 30 |
| May 11^th^ - 20^th^ | 1,885 | 30 |
| May 21^th^ - 31^st^ | 987 | 30 |
| Jun 1^st^ - 10^th^ | 681 | 30 |
| Jun 11^th^ - 20^th^ | 457 | 30 |
| Jun 21^st^ – 30^th^ | 279 | 30 |
| Jul 1^st^ – 10^th^ | 133 | 30 |
| Jul 11^th^ - 20^th^ | 501 | 30 |
| Jul 21^st^ - 31^st^ | 603 | 30 |
| Aug 1^st^ – 10^th^ | 1,238 | 30 |
| Aug 11^th^ - 20^th^ | 1,707 | 30 |
| Aug 21^st^ - 31^st^ | 874 | 30 |
| Sep 1^st^ – 10^th^ | 299 | 30 |
| Sep 11^th^ - 20^th^ | 18 | 18 |
| Total | 40,527 | 618 |

The genetic strains were obtained from Global Initiative on Sharing all Influenza Data (GISAID), accessed via https://www.gisaid.org/.

### **S2. Quantifying the time-varying genetic distance**

As one of the first published sequences from China, Wuhan-Hu-1/2019|EPI_ISL_402125, which was collected in December 31^st^, was set as an initial strain in this study. We calculated the pair-wise genetic distance based on the initial strain. Let *t* denotes the time period (or date), and for the *i*-th virus sample, the amino acid sequence can be denoted as

$X_{i}^{t}=\{x_{ij}^{t}\}$, *j* = 1, …, *J*, and *i* =1, …, *n_t_*

where index *j* indicates the *j*-th codon position in the sequence, and *n_t_* is the total number of samples in *t*-th period. Let the initial strain be $V=\{v_{ij}\}$, and the time-varying average genetic distance $d^{t}$ to the initial strain is defined by the Hamming distance as follows,

$d^{t}=\sum_{i=1}^{n_{t}} \sum_{j=1}^{J} I(v_{ij}\neq x_{ij}^{t})/n_{t}$.

The genetic distance from the initial strain is supposed to gradually increase if there is an accumulation of mutation with evolutionary advantage. To better observe the dynamic trend of genetic distance, a sliding window was applied to the whole study periods. Let *W* denotes the window size that indicates a constant time period (e.g., one week or half a month), and *s* denotes the step length that indicates the distance that the window slides each time. Hence, for $d^{t}$ in time *t*, we calculate the average genetic distance based on sample strains collected from *t*-*W*/2 to *t*+*W*/2. And for the next time *t*+1, sample strains collected from *t*-*W*/2+*s* to *t*+*W*/2+*s* were included in calculation. In this study, the window size *W* was fixed to be 14 (days), and step length was 2 (days).

The statistical analysis was conducted by ***R*** (version 3.6.3).

### **S3. Time-varying genetic distance in ORF1**

A set of non-structural proteins (nsp) produced as cleavage products of the ORF1 viral polyproteins assemble to facilitate viral replication and transcription. And the time-varying genetic distances in these proteins were shown as below. The continuous increasing trends of genetic distance were found in the nsp3 and the RdRp region, while in other proteins, the genetic distance always fluctuated below 0.5 codon (the red dash line) and did not show a clear trend.

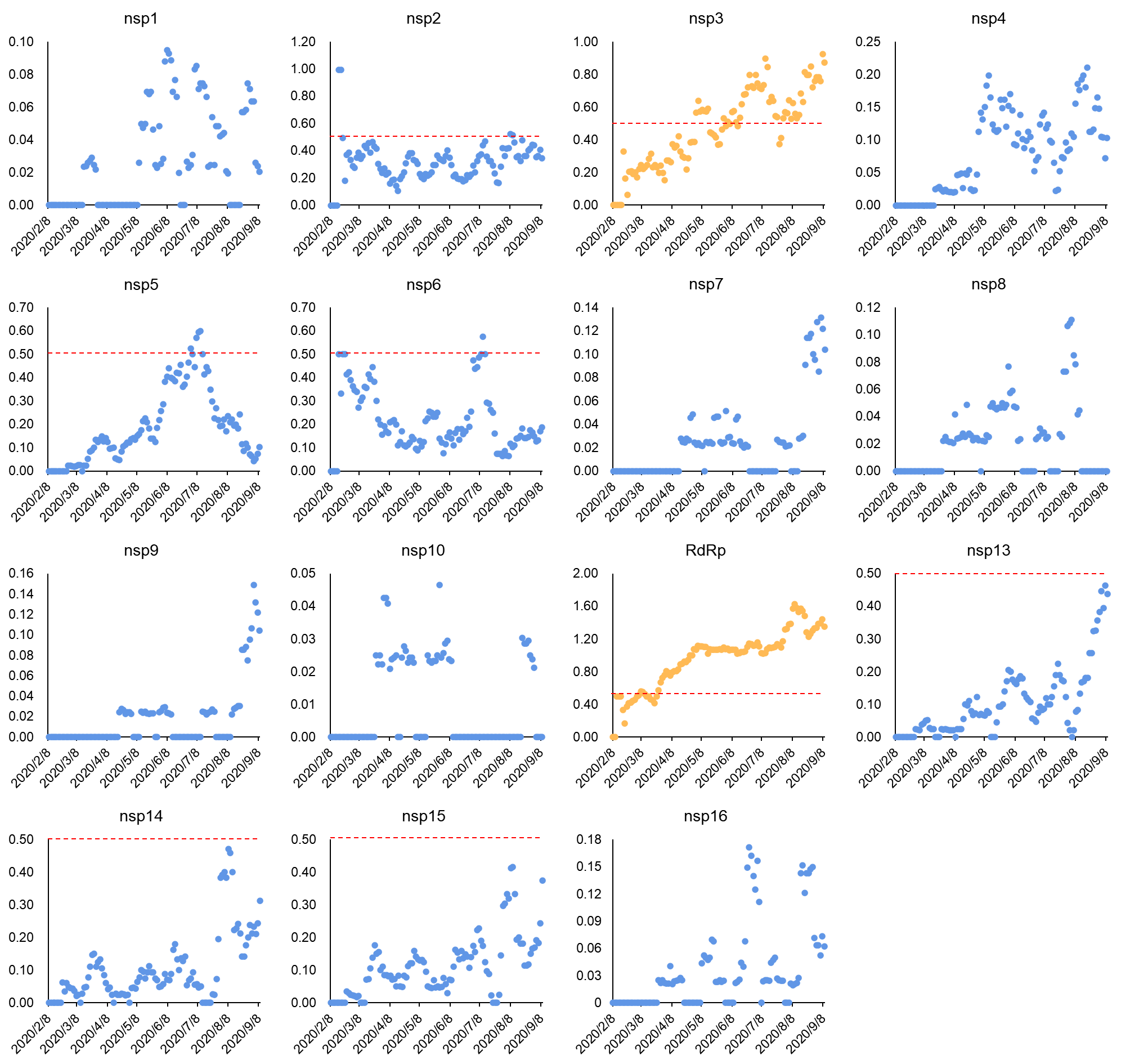

Figure S3. Time-varying genetic distance in ORF1.

The yellow dots represent genetic distance in nsp3 and RdRp with continuous increasing trends. And the blue dots represent distance in other non-structure proteins with weak trends. The red dash line indicates genetic distance equaling to 0.5 codon.
